# *Cauliflower mosaic virus* disease spectrum uncovers novel susceptibility factor NCED9 in *Arabidopsis thaliana*

**DOI:** 10.1101/2022.12.09.519780

**Authors:** Gesa Hoffmann, Aayushi Shukla, Silvia López-González, Anders Hafrén

## Abstract

Viruses are intimately linked with their hosts and especially dependent on gene-for-gene interactions to establish successful infections. The genotype of their hosts thus has a strong influence on the outcome virus disease. On the host side, defence mechanisms like tolerance and resistance can occur within the same species leading to differing virus accumulation in relation to symptomology and plant fitness. The identification of novel resistance genes and susceptibility factors against viruses is an important part in understanding viral pathogenesis and securing food production. The model plant *Arabidopsis thaliana* displays a wide symptom spectrum in response to RNA virus infections and unbiased genome-wide association studies have proven a powerful tool to identify novel disease-genes. In this study we infected natural accessions of *Arabidopsis thaliana* with the pararetrovirus *Cauliflower mosaic virus* to study the phenotypic variations between accessions and their correlation with virus accumulation. Through genome-wide association mapping of viral accumulation differences, we identified several susceptibility factors for CaMV, the strongest of which was the abscisic acid synthesis gene *NCED9*. Further experiments confirmed the importance of abscisic acid homeostasis and its disruption for CaMV disease.

## INTRODUCTION

Plant viruses are ubiquitous in wild and cultivated habitats with profound impacts on host populations (Prendeville *et al*., 2012). As obligate intracellular parasites, they are fully dependent on host-compatibility to complete their replication cycle, and genetic variation within both the plant and viral species can have major effects on the disease outcome (Butkovic *et al*., 2022; Cecchini *et al*., 1998). Of particular interest is the continuum of two mechanisms, tolerance and resistance, plants employ against invading pathogens. Host resistance leads to reduced or absent viral replication and functions commonly through targeted degradation of viral components and incompatibility with the host machinery (Soosaar *et al*., 2005). Tolerance is fundamentally different from resistance and defined as a mitigation strategy aimed at minimizing cost of infection on plant growth, yield and reproduction, rather than investing resources to fight the infection by suppressing pathogen multiplication (Pagán & García-Arenal, 2020; Cooper & Jones, 1983). Tolerance mechanisms can result in high levels of virus accumulation in visible healthy plants, a powerful example being the recently reported *Arabidopsis latent virus 1* that has spread through natural and laboratory populations of *Arabidopsis thaliana* (Arabidopsis) without detection (Verhoeven *et al*., 2022). While agricultural research has historically focused on resistance to battle virus disease, evidence is accumulating that tolerance plays a pivotal role for many plant-virus interactions, especially in natural ecosystems, where most plants are infected by at least one virus at any given time but still appear healthy (Paudel & Sanfaçon, 2018; Roossinck, 2013). Identifying the underlying genetics of the tolerance-resistance spectrum is a difficult task, with genome-wide association studies (GWAS) emerging as a potential tool to find novel genes and pathways implicated in plant-pathogen interactions (reviewed in (Bartoli & Roux, 2017)). Compared to other pathogen classes, GWAS on plant-virus interactions are scarce and most have focused on crop and vegetable species (reviewed in (Monnot *et al*., 2021)). Even though, thanks to the extensive 1001 genomes project, Arabidopsis is a superb resource for GWA studies, with over 1000 sequenced naturally inbred accessions collected worldwide (Consortium, 2016). To our knowledge six recent GWA studies have been conducted on RNA virus infections in Arabidopsis (Butkovic *et al*., 2022; Liu *et al*., 2022; Butković *et al*., 2021; Montes *et al*., 2021; Rubio *et al*., 2019; Pagny *et al*., 2012) and successfully identified genetic loci impacting viral infections.

In addition to discovering new disease and resistance genes for possible application in crop breeding and protection strategies, natural genetic variation and associated phenotypic variation in virus accumulation and symptomology can suggest fundamental perspectives on plant-virus interactions. Only one of the six virus/Arabidopsis GWA studies determined both symptomology and virus accumulation and found a weak positive correlation between the traits (Rubio *et al*., 2019). Yet, plant viruses generally do not show an correlation between symptomology and accumulation across Arabidopsis accessions as observed e.g. for CaMV and other viruses in more narrow experimental setups (Bergès *et al*., 2021; Shukla *et al*., 2018; Pagán *et al*., 2007; Cecchini *et al*., 1998), suggesting tolerance as a ubiquitous process in plant viral diseases.

In this study we examined the disease spectrum of the double-stranded DNA Caulimovirus *Cauliflower mosaic virus* (CaMV; family *Caulimoviridae*) in 100 natural accessions of *Arabidopsis thaliana*. CaMV host range is limited to members of the Brassicaceae, including mustard, broccoli and cabbage and infects natural populations of Arabidopsis (Pagán *et al*., 2010). CaMV challenges its host with the establishment of large cytoplasmic viral replication centers, as well as an uncommon increase of global translation, due to the viral translational transactivator protein P6 (Hoffmann *et al*., 2022; Schoelz & Leisner, 2017). The unique properties of CaMV implicate the existence of a network of host factors possibly influencing CaMV disease. Interestingly, CaMV infection was shown to cause a range of disease severity in response to water deficit in natural accessions of Arabidopsis (Bergès *et al*., 2020), altogether making CaMV a suitable virus for a GWA study in Arabidopsis.

Here, we show that CaMV disease differs greatly in Arabidopsis accessions, dependent on the host genotype and use this variety to map underlying host genes. We find that the abscisic acid (ABA) synthesis gene 9-Cis-Epoxycarotenoid Dioxygenase 9 (*NCED9*) is an important susceptibility factor for CaMV, as infection is almost completely abolished in the *nced9* mutant line. Additionally, ABA, an important plant hormone in abiotic and biotic stress response (Verma *et al*., 2016; Ton *et al*., 2009), is targeted during CaMV infection and miss-regulation of ABA homeostasis increases CaMV levels.

## Material & Methods

### Plant Material and Growth Conditions

100 accessions of *Arabidopsis thaliana* (supplementary Table S1) were provided by the group of Magnus Nordborg (Gregor Mendel Institute, Vienna). The T-DNA lines used in this study were ordered from the NASC stock center and all generated in Columbia (Col-0) background, which was taken as control for all mutant experiments (supplementary Table S2). Seeds were plated on damp soil and stored at 4C in the dark for one week to ensure germination synchronization. Seedlings were separated at 6 plants per pot 8 days after transfer to a walk-in chamber in short day conditions (120 mE, 10h light / 14h dark cycle) at 22°C and 65% relative humidity. Pots were randomized within each tray and tray position within the chamber was switched randomly once a week. Infections were carried out 18 days after transfer to growth conditions. Infections of natural accessions were repeated twice in timely separated experiments. T-DNA lines were infected at least three times in timely separated experiments. Arabidopsis plants were grown in walk-in chambers in standard long day conditions (16h light / 8h dark cycle) at 22°C and 65% relative humidity for propagation. For long day infection experiments, seeds were plated on damp soil and stored at 4C in the dark for one week to ensure germination synchronization. Seedlings were separated at 4 plants per pot 6 days after transfer to a walk-in chamber and infections carried out 15 days after transfer to growth condition.

### Virus Inoculation and Symptom Scoring

Arabidopsis plants were infected with CaMV 18 days after germination. The first true leaves were infiltrated with *Agrobacterium tumefaciens* strain C58C1 carrying CaMV strain CM1841. Plants were scored for symptoms and photographed at 21 dpi. Symptom categories (0-5) correspond to no visible symptoms (0), mild vein clearing (1), leaf bending (2), rosette distortion (3), rosette shrinking (4) and early senescence with necrotic lesions (5) and were determined for each accession. Failed infections were removed from pots before taking above ground fresh weights for individual plants. All infected plants (n=3-6) of one accession were pooled for titer measurements and ground to fine powder in liquid nitrogen. Infections with the RNA viruses were performed using clones described in (Ling *et al*., 2013) for *Turnip rosette virus* (TRoV – family *Solemoviridae*) and (Garcia-Ruiz *et al*., 2010) for and Turnip mosaic virus (TuMV – family *Potyviridae*).

### Virus Quantification and gene expression analysis

For CaMV DNA quantification, 100mg pulverized frozen leaf material was resuspended in 300 μl 100mM Tris buffer (pH 7.5), supplemented with 2% SDS and treated with Proteinase K. Total DNA was precipitated with isopropanol 1:1 (v:v). RNA extraction from rosette tissue was performed with a Qiagen RNeasy kit and on-column Dnase I digestion according to the manufacturer’s protocol. About 500 ng of total RNA was used for first-strand cDNA synthesis with a Maxima First Strand cDNA Synthesis Kit (Thermo Fisher Scientific Waltham, MA, USA]). qRT-PCR analysis of DNA and cDNA was performed with Maxima SYBR Green/Fluorescein qRT-PCR Master Mix (Thermo Fisher Scientific) using the CFX Connect Real-Time PCR detection system (Bio-Rad, Hercules, CA, USA) with specific primers (supplementary Table S9). Viral DNA was normalized to genomic *ACTIN7* (AT5G09810) for all accessions and *18S* ribosomal DNA for T-DNA lines. Viral transcripts and ABA responsive transcripts were normalized to *PP2a* (*AT1G69960*).

### Genome-wide association mapping

Genome-wide association mapping was performed on 100 accessions using an online portal provided by Gregor Mendel Institute, Austria (https://gwas.gmi.oeaw.ac.at) (Seren *et al*., 2012) against the Imputed Fullsequence Dataset (Long *et al*., 2013; Cao *et al*., 2011; Gan *et al*., 2011) with an accelerated mixed model to correct for population structure (Seren *et al*., 2012). Analysis was performed with untransformed data. For this publication, SNPs were considered when they withstood a 5% false discovery rate by Benjamini-Hochberg-Yekultieli thresholding (Benjamini & Hochberg, 1995) and a minor-allele count of >=5. 15 T-DNA lines were chosen for the highest scoring SNPs that fell into gene bodies, caused a miss-sense mutations and had available T-DNA insertions in the NASC stock centre.

### Chemical Treatments

For chemical treatments, abscisic acid (Sigma-Aldrich, A1049) and Nordihydroguaiaretic acid (Merck Chemicals and Life Science, 74540) were prepared in 99% EtOH for stock solutions. 17-day-old seedlings were sprayed with dilutions 24h before infection. The treatment was repeated once a week at the same time until harvest. The last application was performed 24h before harvest.

### Broad-sense heritability calculation

The estimation of broad-sense heritability (h2b) was calculated as the percentage of the total variance accounted by genetic (accession) differences (h2b = σ2 G/σ2 P, were σ2 G is the genetic variance component of σ2 P total phenotypic variance). σ2 P and σ2 G were derived by variance components analysis using separated univariate analyses (Shukla *et al*., 2018).

### Transcriptome analysis

Transcriptome data were generated by Chesnais et al. 2022. For the re-analysis of the bulk RNA-seq data, raw data was downloaded from BioProject number PRJEB49403 from the European Nucleotide Archive (https://www.ebi.ac.uk/ena/browser/view/PRJEB49403). Analysis was done on three replicates of mock and CaMV infected samples. In brief, downloaded reads were trimmed and checked with TrimGalore (Version 0.5.0; https://github.com/FelixKrueger/TrimGalore, based on Cutadapt (Martin, 2011)) using the options -q 20 --fastq --stringency 1 --length 32 --paired. Afterwards reads were mapped to the TAIR10 genome using Tophat2 (Version 2.1.1; (Kim *et al*., 2013)) with the parameters --library-type=fr-firststrand -g 1 -a 10 -i 40 -I 5000 -r 150 using a the TAIR10 reference annotations for all annotated genes. Mapped output files were sorted and indexed using samtools (Version 1.6; (Li *et al*., 2009)). FeatureCounts from the subread package (Version 2.0.1; (Liao *et al*., 2014)) was used with the options -T 8 -p -t gene -O -s 2 against all genes in the TAIR10 genome to generate a counts table for subsequent analysis differentially expressed genes using the R package Deseq2 ((Love *et al*., 2014)).

### Bioinformatics

Plots were made with R 4.0.2, using the packages “ggplot2” (Wickham, 2016), “tidyverse” (Wickham *et al*.), “raincloudplot” (Allen *et al*., 2019) or base functions. All statistical calculations were performed in R with base functions. Figure arrangements were finalized using AffinityDesigner 1.10. Latitude and longitude data, as well as SNP data and impact prediction were taken from the https://1001genomes.org/ website and the POLYMORPH1001 tool (https://tools.1001genomes.org/polymorph/index.html) (supplementary Table S1). Gene loci and descriptors were assembled through the PANTHERDB website version 16.0 using bedtools v2.30.0 “closest” function.

## RESULTS

### CaMV disease severity is highly variable in Arabidopsis

CaMV, like Arabidopsis is distributed worldwide and infects Arabidopsis and other Brassicaceae in wild populations (Pagán *et al*., 2010; Raybould *et al*., 1999). In this study, we examined CaMV disease in 100 Arabidopsis accessions under controlled conditions. Accessions exhibited a broad range of symptoms that were scored at 21 days past infection (dpi). We categorized symptoms from mild vein clearing (1) and leaf bending (2) over rosette distortion (3) and rosette shrinking (4) to early senescence with necrotic lesions (5) (Figure 1A; supplementary Figure S1A). Only two accessions, PHW-3 and IP-Oja-0, did not develop any visible disease (0), while most accessions developed moderate symptoms (supplementary Table S3). 83 of the 100 tested accessions were collected in Europe (Figure 1B, supplementary Figure S1B for world map), but we could not find clustering of similar disease severities along the longitudinal or latitudinal gradient (Figure 1B) and the main Admixture groups (n>5) in our dataset did not reveal a pattern in symptom severity (Figure 1C). Relative fresh weight after virus infection is a widely used proxy for disease severity and it was strongly correlated with the visually determined disease categories in our dataset (Figure 1D, supplementary Table S3). Importantly, virus induced fresh weight loss did not correlate with the total fresh weight of mock inoculated plants, indicating that virus disease costs in these conditions are not dependent on differences in growth capacity between individual accessions (supplementary Figure S1B). We also tested a few accessions under different light conditions to evaluate the robustness of accession-specific symptomology and found that the range of symptoms was essentially reproduced (compare Figure 1A and supplementary Figure S1C). These results reveal a large spectrum of CaMV induced symptoms in Arabidopsis that appears largely independent of the global origin of the accession when grown in controlled conditions.

**Figure 1:**
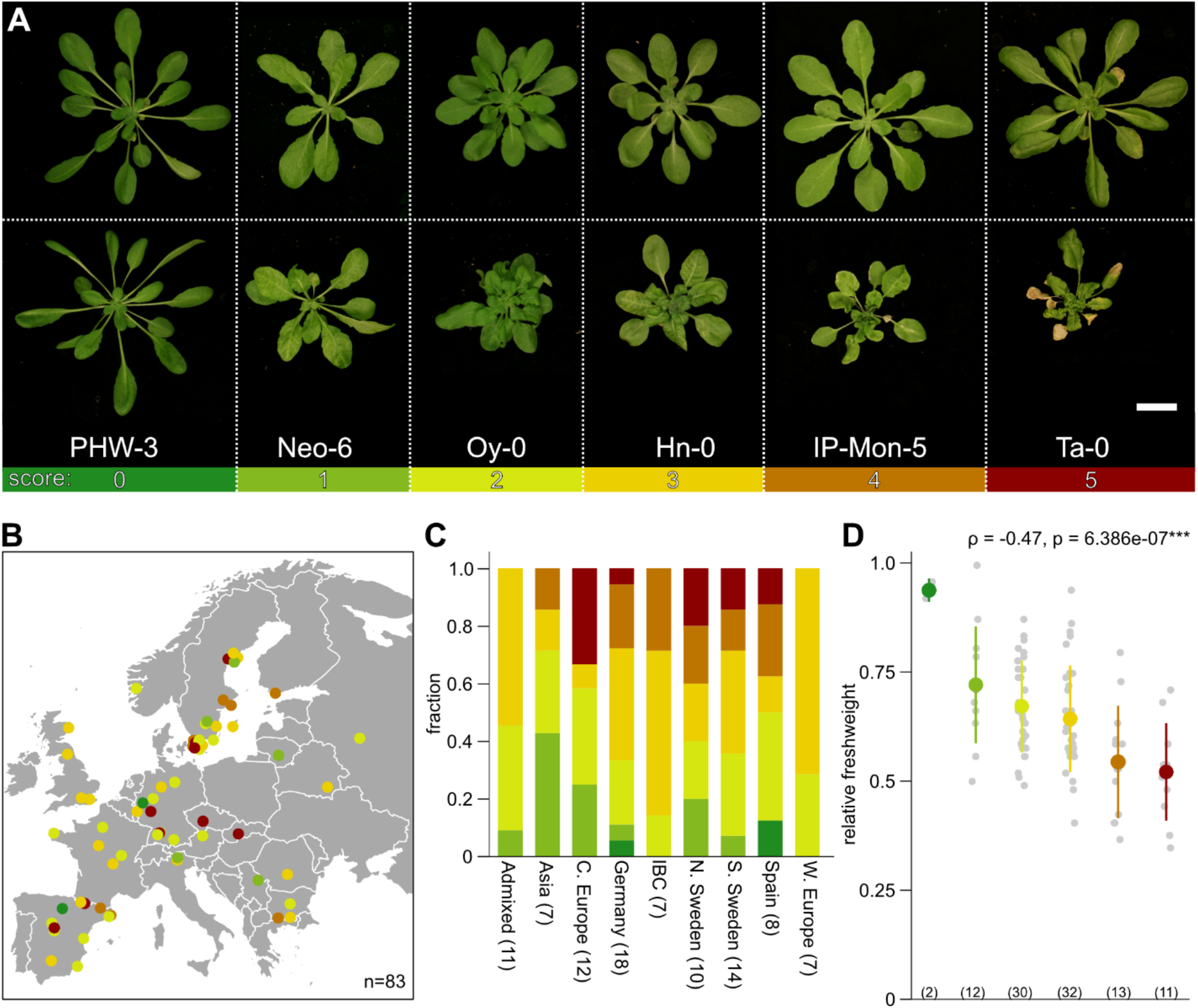
The broad spectrum of CaMV disease in Arabidopsis. **(A)** Representative images of symptom range induced by CaMV infection 21 dpi. Upper panel: mock infected plants, lower panel: CM1841 infected plants. Accession identifier is written below. Colors correspond to symptom categories. Scale bar = 2 cm **(B)** Geographical distribution of 83 Arabidopsis accessions from Europe, representing 83% of examined accessions. Dot colors indicate symptom categories. **(C)** Fraction of symptom categories divided by Admixture groups. Number of accessions in each admixture group is indicated in brackets. (IBC = Italy_Balkan_Caucasus C = Central, N = North, S = South, W = Western) **(D)** Dot blot of relative fresh weight of accessions in symptoms categories. (n) indicate numbers of accessions in each category. Coloured dot and stick represent mean ± standard deviation. Grey dots represent individual accessions. Correlation was calculated with Spearman rank test.

### Tolerance and resistance govern CaMV disease in Arabidopsis

To evaluate the relation between virus accumulation and disease symptomology, we determined viral genomic DNA levels in parallel with the symptom scoring and fresh weight analysis presented in Figure 1. CaMV DNA accumulation was measured from pools of infected plants from the two replicate experiments with good reproducibility (Figure 2A, supplementary Table S4). We detected a 28-fold difference between the highest virus DNA measurement (IP-Ven-0) and the lowest (Lerik1-4) in symptomatic plants. Interestingly, we found only a weak correlation between viral titer and plant symptoms and the four highest CaMV accumulators belonged to symptom group 1,2,3 and 5, indicating that virus multiplication and virulence are largely uncoupled in the present settings (Figure 2B). Likewise, several accessions from the severe symptom categories (4 and 5) accumulated low levels of virus, suggesting hypersensitivity. A equally poor, but positive correlation between symptoms and viral accumulation has been previously described for the potyvirus *Turnip Mosaic Virus* (TuMV) (Rubio *et al*., 2019), while no correlation was found for cucumovirus *Cucumber Mosaic Virus* (CMV) (Pagán *et al*., 2007), altogether strengthening that disease symptoms are frequently not a consequence of the amount of virus within a plant. We also could not detect differences in CaMV accumulation between the different admixture groups (Figure 2C), except a slightly higher value for relics, but the low number of accessions in this group might confound the effect. Again, the highest CaMV accumulators were scattered between the Admixture groups. These data show that many Arabidopsis accessions vary in their tolerance against CaMV in a manner largely uncoupled form accumulation and thus, that symptom development in individual accessions is far from a direct indicator for CaMV accumulation (Figure 2D).

**Figure 2:**
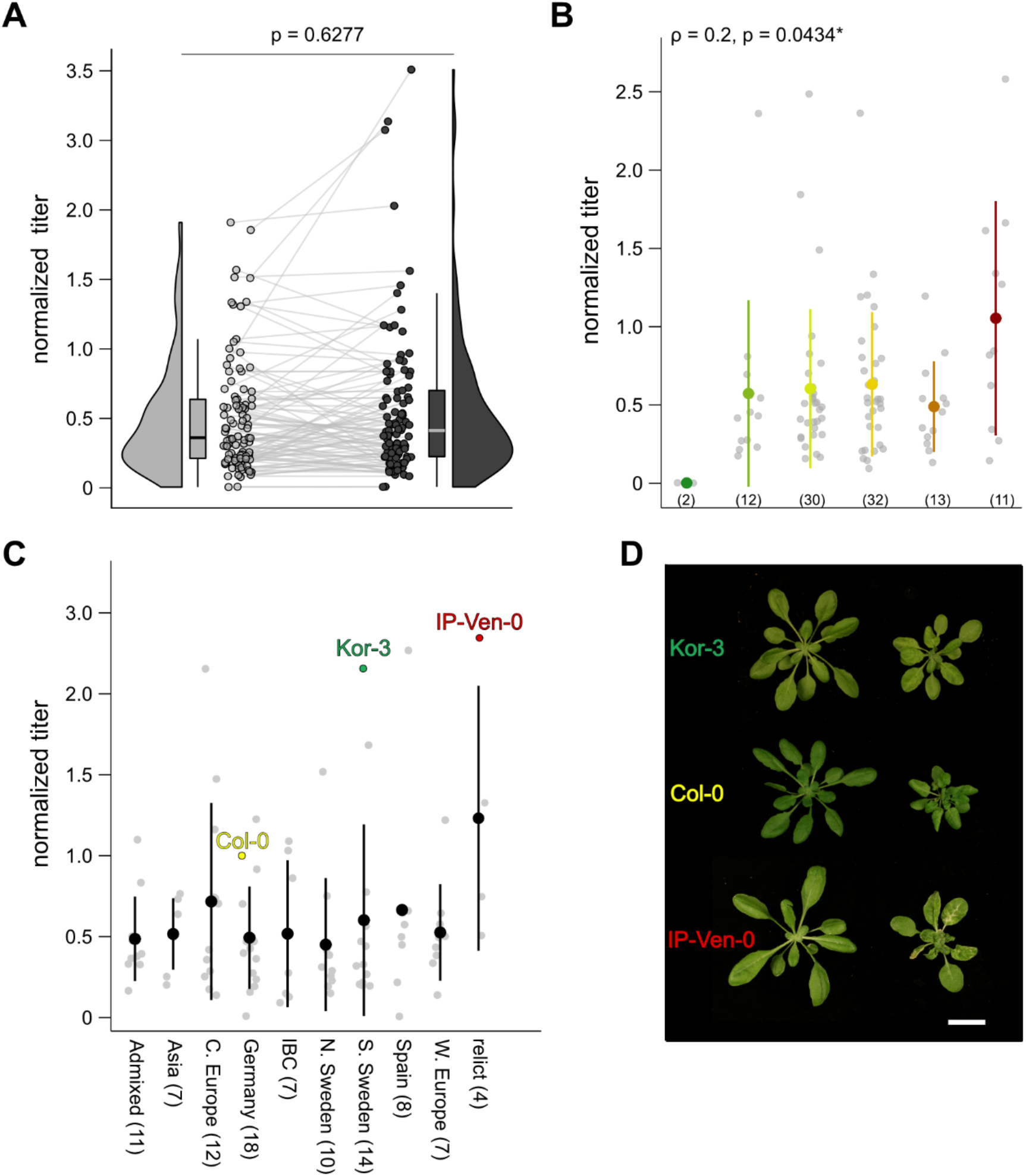
CaMV accumulation only weakly correlates with Arabidopsis symptoms. **(A)** Raincloud plot of CaMV DNA accumulation in 100 accession 21 dpi in two independent replicates. P-value was calculated by Kruskal-Wallis rank sum test. **(B)** Dot blot of CaMV DNA accumulation between symptoms categories. (n) indicate numbers of accessions in each category. Colour corresponds to symptom categories (Figure 1A). Coloured dot and stick represent mean ± standard deviation. Grey dots represent individual accessions. **(C)** Dot blot of CaMV DNA accumulation between Admixture groups. Accessions depicted in (D) are highlighted. Number of accessions in each admixture group is indicated in brackets. Black dot and stick represent mean ± standard deviation. Grey dots represent individual accessions. **(D)** Representative image of accessions xxx and Col-0. Both accessions accumulate twice as much CaMV DNA as Col-0 but fall on either side of Col-0 on the disease spectrum. Scale bar = 2 cm

A recent study by (Liu *et al*., 2022) examined the quantitative resistance of Arabidopsis against two distantly related strains of CMV. 41 (for CMV-Q) and 42 (Fny-CMV-Δ2b) accessions were shared between their study and ours. Interestingly, while no correlation could be detected between CaMV and CMV accumulation in general, individual accessions like IP-Ven-0 accumulated high virus loads in both cases and IP-Oja-0 showed full resistance against CaMV and accumulated very low levels of CMV (supplementary Figure S2). Another study on CMV virulence in Arabidopsis accessions from the Iberian peninsula, also found that CMV infection in IP-Ven-0 drastically reduced seed production (−96.5%), while IP-Oja-0 seed production was only decreased by −20% after infection (Montes *et al*., 2021). The absence of a global correlation between CaMV and CMV accumulation over the accessions suggests that individual plant-virus interactions are commonly of high importance, but single accessions might still exhibit strong resistance or susceptibility to viruses generally, possibly as a consequence of physiological traits.

### Genome-wide association mapping identifies novel CaMV susceptibility factors

We used the GWAPP tool (Seren *et al*., 2012) to conduct a genome-wide association (GWA) mapping of symptoms, relative fresh weight and relative CaMV accumulation in the 100 accessions. It is important to note that 100 accessions are a small sample size for GWA-mapping, which will result in limited resolution. Neither symptom-category, nor relative fresh weight data resulted in the identification of SNPs above the Benjamini-Hochberg threshold (supplementary Figure S3), however several regions were associated with CaMV accumulation (Figure 3A, supplementary Table S5). Broad-sense heritability for CaMV DNA accumulation was 0.58, similar to previous observations in plant-virus systems (Monnot *et al*., 2021; Shukla *et al*., 2018). After thresholding, we found 140 genes within a 2 kb region of significant SNPs for CaMV titer (supplementary Table S6), in accordance with the multifaceted process of viral replication. Most associated genes have no annotated function in ThaleMine (v.5.1.0-20221003). A protein class ontology search on PantherDB.org (v17.0) showed that the largest group of genes (16) by protein class ontology encodes for metabolite interconversion enzymes (PC00264), eight of which are oxidoreductases (PC00176), followed by protein modifying enzymes (8; PC00260) and transcriptional regulators (6, PC00264). Since viral replication and accumulation could be influenced by yet unknown mechanisms, we did not want to limit our downstream analysis and randomly selected 15 SNPs above the threshold located in gene bodies that caused miss-sense mutations (Coloured arrowheads in Figure 3A; supplementary Table S2) for which we analysed CaMV accumulation in Col-0 based T-DNA insertion lines.

**Figure 3:**
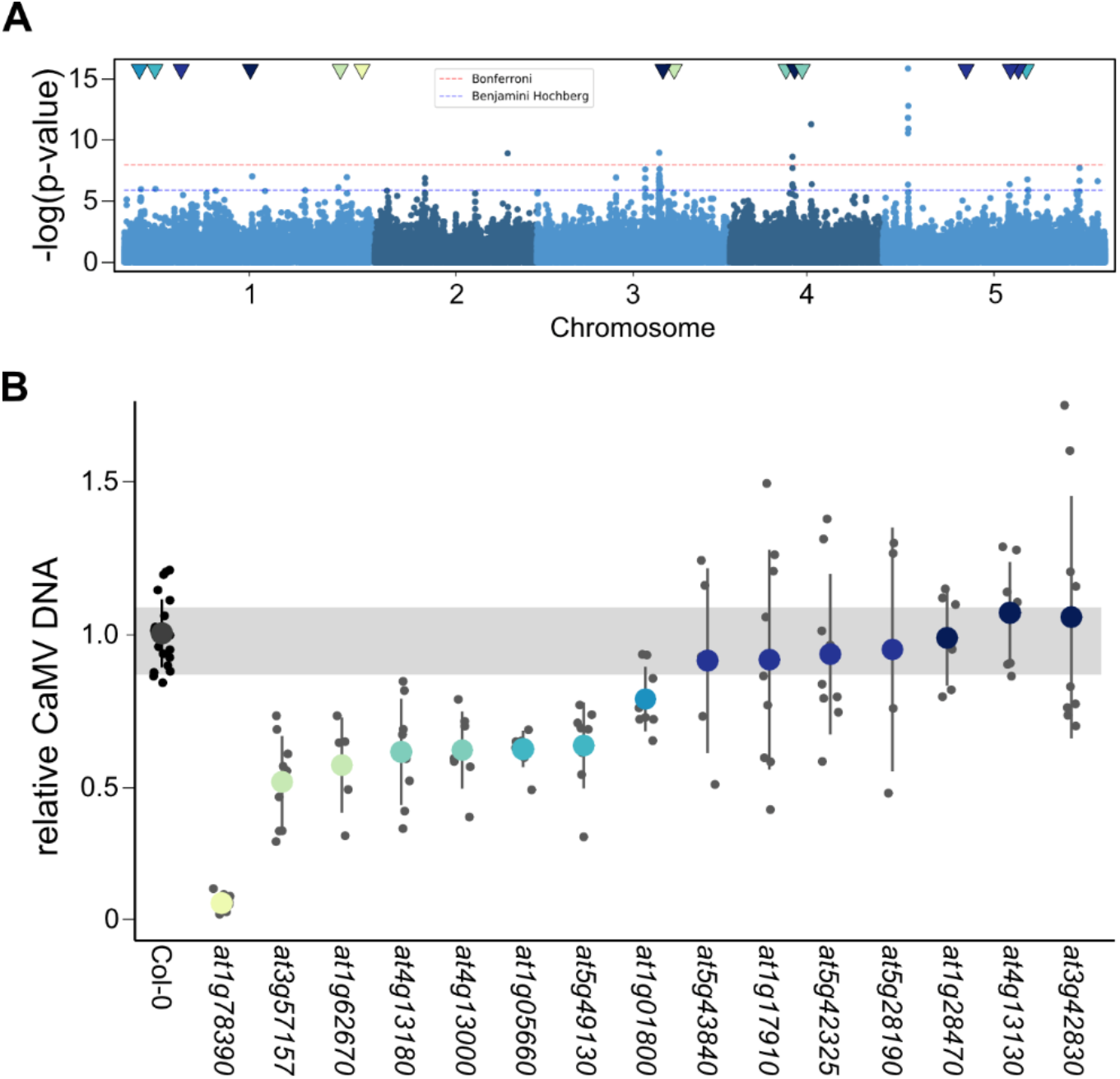
Genome-wide association mapping of CaMV accumulation and candidate screening. **(A)** Manhattan plot of GWA results for CaMV accumulation in 100 natural accessions. Blue shading corresponds to the five Arabidopsis chromosomes. Blue lines indicate significance threshold after Benjamini-Hochberg correction, red line represents the more stringent Bonferroni multiple testing correction. **(B)** Relative CaMV DNA accumulation in T-DNA lines of GWA candidates (indicated by ATG-number) at 21 dpi compared to Col-0 wild type. n = 4-22. T-DNA lines are listed in supplementary dataset2 Table S2. Colored dot and stick represent mean ± standard deviation. Grey dots represent individual accessions. Grey bar represents standard deviation of Col-0.

Intriguingly, of the 15 tested lines eight showed a significant reduction in CaMV accumulation compared to Col-0 (Figure 3B). It is noteworthy that none of the tested lines increased CaMV accumulation, suggesting that our GWA mapping mainly identified susceptibility factors. All lines developed Col-0 like symptoms at 21 dpi, except for SALK_123975.34.85.x, which also had the most striking reduction of viral DNA (~5% Col-0). This line harbors an insertion in the only exon of AT1G78390 (Lefebvre *et al*., 2006). AT1G78390 encodes for 9-Cis-Epoxycarotenoid Dioxygenase 9 (NCED9), an enzyme involved in the biosynthesis of abscisic acid (ABA). The identified SNP causes a missense mutation of Valin415 to Leucin in the NCED9 coding sequence, with a predicted moderate effect (Figure 4A). This particular polymorphism only occurs in 29 natural accessions, that are all but one clustered in central and northern Europe and Russia (FIGURE 4B). We expanded the virus accumulation analysis and additionally tested 5 additional lines harboring this SNP (supplementary Table S7). On average, accessions with NCED9-415L accumulated significantly more virus than NCED9-415V (Figure 4C).

**Figure 4:**
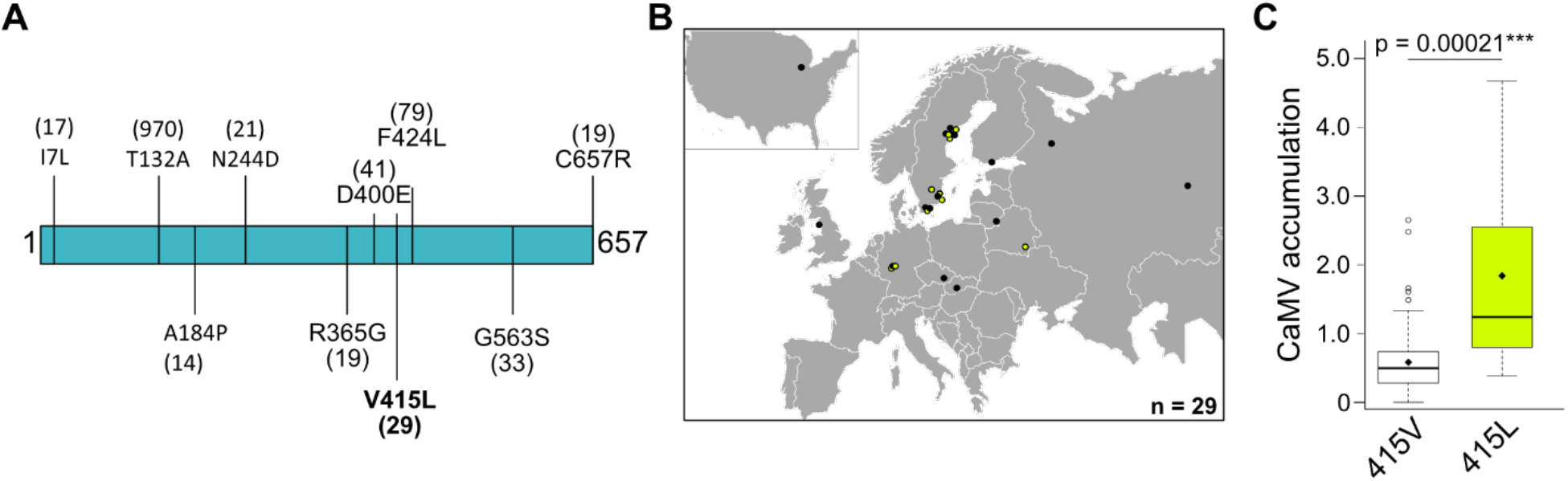
Allelic variation in NCED9 influences CaMV accumulation. **(A)** Graphic representation of NCED9 protein (657 AA) with AA substitutions due to SNPs present in more than 10 accessions annotated from POLYMORPH 1001 browser. **(B)** Geographical distribution of 29 Arabidopsis accessions harbouring NCED9-415L. Yellow dots indicate accessions in our collection used for CaMV experiments. **(C)** CaMV DNA accumulation relative to Col-0 in NCED9-415V accessions (n=95) and NCED9-415L (n=10) accessions. P-value was calculated using pairwise Wilcoxon rank rum test with continuity correction.

### NCED9 is essential for robust CaMV accumulation

NCED9 is best examined for its role during seed development and germination (Lefebvre *et al*., 2006; Tan *et al*., 2003a). We found that CaMV infection induced *NCED9* expression when compared to healthy plants, albeit still to low levels (Figure 5A). We used an independent publicly available transcriptome set of Arabidopsis infected with the same CaMV strain CM184I from 21 days after aphid inoculation (Chesnais *et al*., 2022) and could also find increased levels of *NCED9* transcript in response to CaMV (Figure 5B). The *nced9* T-DNA line developed no symptoms except for a mild vein clearing phenotype in older leaves over an infection time of 44 days (Figure 5C) and displayed no fresh weight loss compared to uninfected control plants when challenged with CaMV (Figure 5D). This resistance phenotype was persistent also under long-day light regimes (supplementary Figure S4A). After backcrossing *nced9* into Col-0, we used symptom development to test whether homozygous *nced9* allele is needed for CaMV resistance. Close to 90% of Col-0 plants developed symptoms upon infection, while 0% of homozygous *nced9* plants did. Three independent segregating F2 populations developed Col-like symptoms with a 69-72% frequency, indicating that a homozygous line of *nced9* is required for CaMV resistance (supplementary Figure S4B). Plant resistance to viruses can be specific to the virus species and sometimes even the viral strain (Takahashi *et al*., 2002). The *nced9* mutant is resistant to two strains of CaMV, the milder CM184I and the more virulent Cabb B-JI strain (supplementary Figure 4C), but susceptible to infections with TuMV and *Turnip rosette virus* (TRoV) (supplementary Figure S4D). Thus, NCED9 appears to be a CaMV specific susceptibility factor. CaMV RNAs are very stable and can accumulate to high levels despite reduction in viral DNA (Hoffmann *et al*., 2022). In *nced9*, all three major viral RNA species were reduced, but not as drastic as the viral DNA (Figure 3B and Figure 5E), but still a remarkable inhibition of infection.

**Figure 5:**
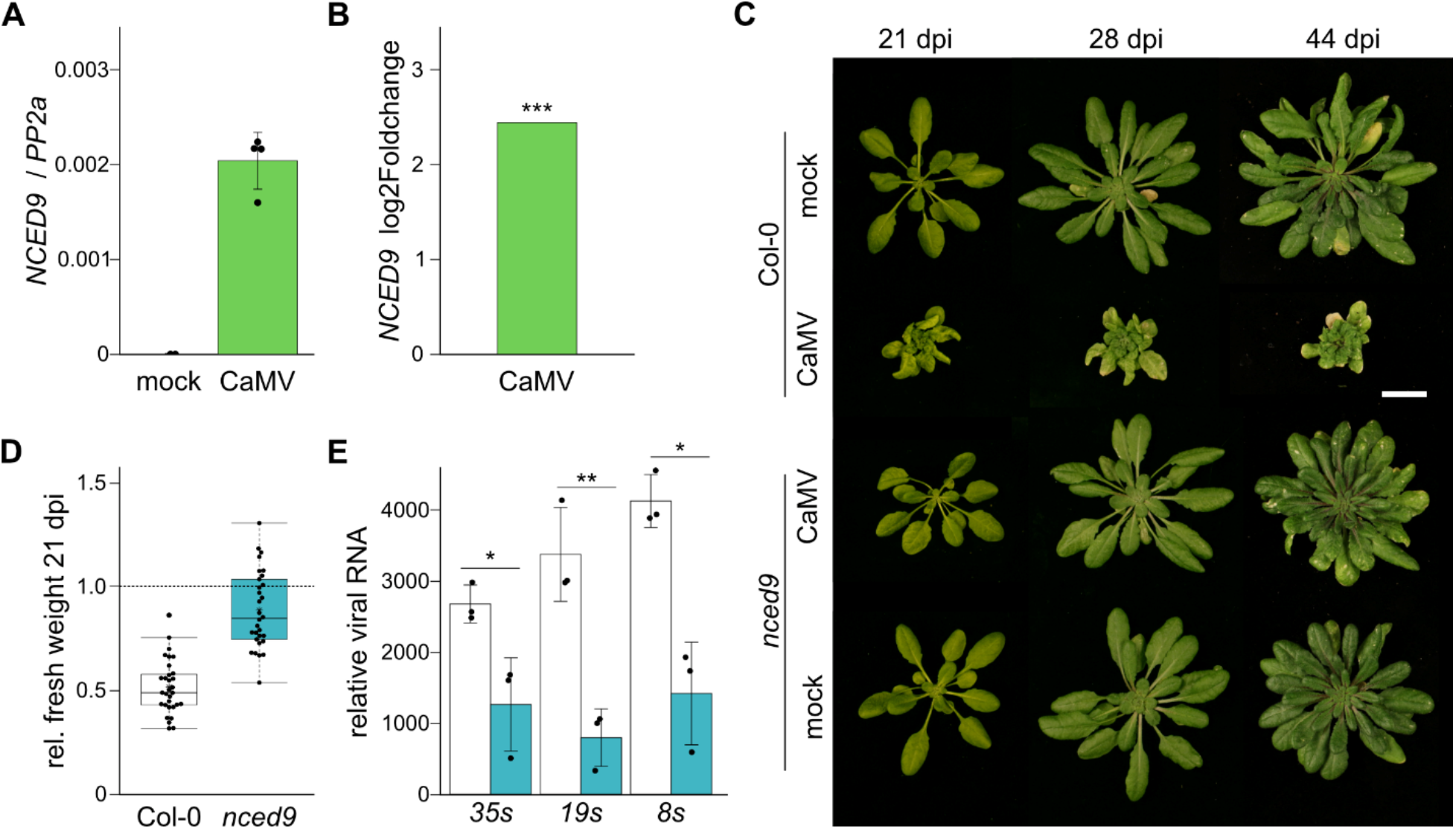
*nced9* is resistant to CaMV infection. **(A)** qRT-PCR of relative transcript accumulation of *NCED9* in mock and CaMV infected Col-0 plants at 21 dpi normalized to PP2a. n=4 **(B)** Log2foldchange of NCED9 in CaMV infected compared to mock plants in the transcriptome dataset of (Chesnais *et al*., 2022) **(C)** Representative image of Col-0 and *nced9* plants after 21, 28 or 44 days after infection with CM184I. Scalebar = 2 cm **(D)** Relative fresh weight of infected Col-0 and *nced9* plants at 21 dpi. Red line indicates mock. **(E)** qRT-PCR of relative transcript accumulation of viral RNAs in Col-0 and *nced9* plants at 21 dpi normalized to PP2a. n=3

### Exogenous ABA application enhances CaMV accumulation in Col-0

The established role of NCED9 in ABA biosynthesis prompted us to investigate the involvement of ABA during CaMV infection. ABA plays multifaceted roles during plant pathogen interactions and exogenous ABA application was found to either increase or reduce pathogen load *in planta* (Alazem *et al*., 2014). We treated seedlings with ABA 24h before infection with CaMV and once a week throughout the 3-week infection time course, the last treatment being 24h before harvest of the whole rosette. Exogenous ABA spray application reduced Col-0 rosette growth in a concentration dependent manner (Figure 6A & B). The *nced9* plants behaved comparable to Col-0, showing that the line has not lost its sensitivity to ABA (Figure 6A & B). The well-described *aba*2 mutant accumulates about 20-25% of wildtype ABA levels during undisturbed growth (González-Guzmán *et al*., 2002), has a severe growth phenotype and is prone to wilting (Figure 6A). The *aba2* growth phenotype was fully rescued by exogenous ABA spraying, suggesting successful application (Figure 6B). CaMV accumulation in Col-0 was not affected after spraying with low concentrations of ABA (10 or 50 μM), but 100 μM and more strongly 200 μM increased viral DNA content (Figure 6C). Likewise, while virus levels were reduced in non-treated *aba2-1* plants, virus load significantly increased upon 200 μM ABA spray (Figure 6C). Intriguingly, exogenous ABA application had no effect on CaMV accumulation in the *nced9* background, seemingly uncoupling the function of *NCED9* in CaMV infection from bulk ABA synthesis (Figure 6C). The phenolic antioxidant Nordihydroguaiaretic acid (NDGA) is a commonly used inhibitor of lipoxygenases (NCEDs) and as such is an inhibitor of ABA synthesis (Han *et al*., 2004; Creelman *et al*., 1992). NDGA has been used previously in plant-virus studies and either made the plants more susceptible to the virus (He *et al*., 2021) or reduced viral load *in planta* (Alazem *et al*., 2014). We observed that NDGA treatment decreased plant growth in uninfected plants (Figure 6A), but also that NDGA treatment increased CaMV DNA accumulation in Col-0 and *aba2-1*, while it had no effect on virus accumulation in *nced9* (Figure 6D). These results suggests that disturbance of ABA homeostasis, rather than ABA levels might aid virus accumulation. We used CaMV transcriptome data (Chesnais *et al*., 2022) to visualize the effect of CaMV infection on ABA responsive genes in four week old rosettes (Hoth *et al*., 2002). CaMV infection altered the expression of positively and negatively ABA-regulated genes drastically and in an unspecific manner, indicating a disturbance in ABA signaling pathways (Figure 6E and F, supplementary Table S8). To validate that these changes hold true in our experimental conditions, we chose four ABA responsive genes that are downregulated during CaMV infection according to the transcriptomics data and tested their expression with qRT-PCR, indeed confirming their strong transcriptional repression during CaMV infection (Figure 6G). Taken together, our data suggest that CaMV infection benefits from the disturbance of ABA homeostasis, probably through the miss regulation ABA-dependent pathways that ultimately helps viral accumulation. But detailed importance of NCED9 for CaMV as part of these ABA-related mechanisms remains to be determined.

**Figure 6:**
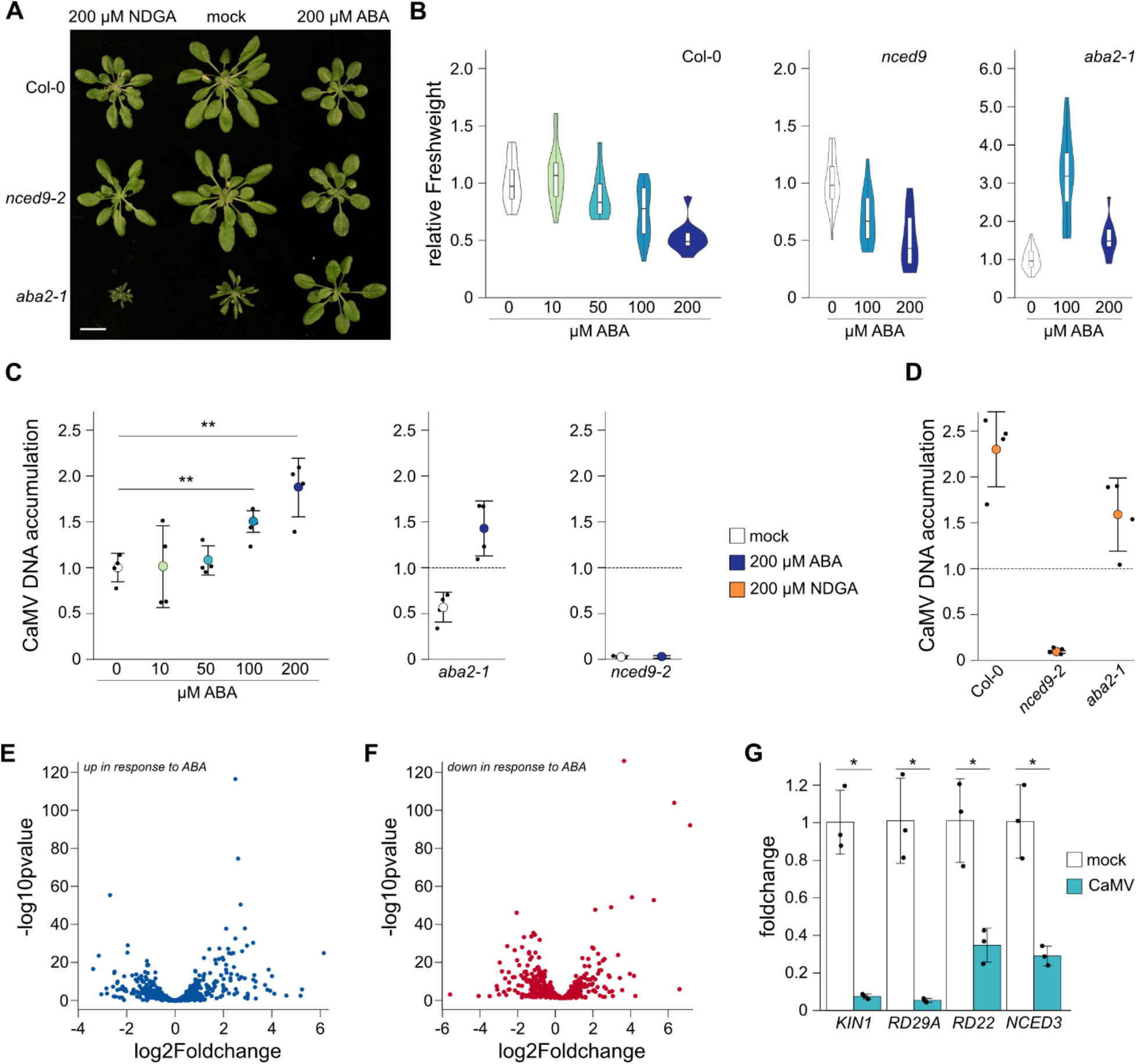
exogenous application ABA enhances CaMV accumulation in a dose dependent manner. **(A)** Representative image of mock inoculated plants after three treatments with either ABA or NDGA. Scalebar = 2 cm. **(B)** Violin plot of relative fresh weight of mock inoculated Col-0 (left panel), *nced9* (middle panel) and *aba2* (right panel) plants after three treatments with indicated concentrations of ABA. **(C)** Relative CaMV DNA accumulation at 21 dpi in Col-0 (left panel), *aba2* (middle panel) and *nced9* (right panel) after three treatments with indicated ABA concentrations n = 4 **(D)** Relative CaMV DNA accumulation at 21 dpi in indicated genotypes after three treatments with 200 μM NDGA n = 4 **(E)** Log2foldchange of ABA responsive genes (“upregulated after ABA treatment”, n=651, (Hoth *et al*., 2002)) in CaMV infected compared to mock plants in the transcriptome dataset of (Chesnais *et al*., 2022) **(F)** Log2foldchange of ABA responsive genes (“downregulated after ABA treatment”, n=680, (Hoth *et al*., 2002)) in CaMV infected compared to mock plants in the transcriptome dataset of (Chesnais *et al*., 2022) **(G)** relative transcript accumulation of ABA responsive genes (from category “up”) in Col-0 plants at 21 dpi mock or CaMV infection normalized to PP2a. n=3

## DISCUSSION

Plants can exhibit amazing plasticity in response to pathogens and the Arabidopsis/CaMV pathosystem is no exception. Arabidopsis is a natural host of CaMV and possibly evolved under CaMV pressure, as proposed for other naturally infecting Arabidopsis viruses (Montes *et al*., 2019). In our conditions, Arabidopsis exhibited a wide spectrum of responses to CaMV that ranged from no symptoms and no viral accumulation to full susceptibility with strong symptoms and high viral accumulation. Notably, we found tolerant and hypersensitive accessions as well, once again exemplifying that symptom severity and virus accumulation are largely uncoupled between host genotypes and that both resistance and tolerance mechanisms shape plant-virus interactions (Figure 7) (Bergès *et al*., 2021; Rubio *et al*., 2019; Pagán *et al*., 2007). The defiance of pathogen-load/symptom connections (“tolerance”) has been reported in other infection-systems, including bacteria and fungi (Gambetta *et al*., 2007; Chen *et al*., 2004), although, examples of clear resistance trajectories also exist, for example for *Pseudomonas syringae* on Arabidopsis that shows strong positive correlation between symptom severity and bacterial density (Kover & Schaal, 2002). CaMV causes moderate symptoms in most accessions, a trend also seen with TuMV in 1050 Arabidopsis accessions (Butkovic *et al*., 2022) and in line with the theory that viruses evolve for an intermediate severity to balance replication and host survival (Torres-Barceló *et al*., 2010). Four accessions (En-2, Wil-2, Sv-0 and Tsu-0) had been previously reported resistant to CaMV infection (Leisner & Howell, 1992). Through our study we have found two additional fully resistant accessions PHW-3 and IP-Oja-0. In accordance, the En-2 resistance locus (CAR1) co-segregates with the microsatellite marker nga128 on chromosome 1 (Callaway *et al*., 1996) and is broken by the P1 protein of CaMV strain NY8153 (Adhab *et al*., 2018), while resistance in Tsu-0 is broken by P6, pointing towards individual resistance mechanisms between the accessions (Hapiak *et al*., 2008).

**Figure 7:**
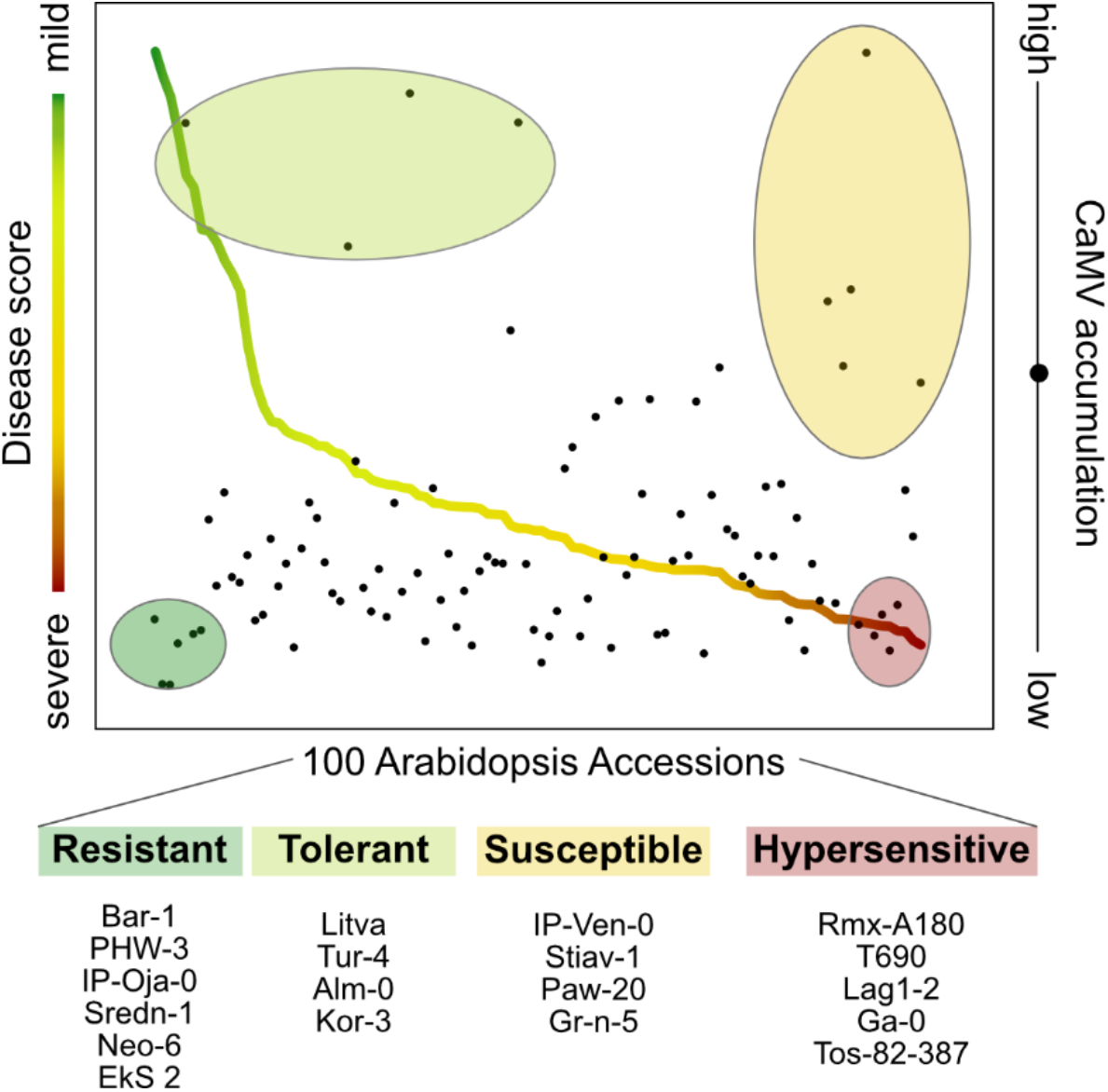
Tolerance and resistance shape CaMV disease in Arabidopsis thaliana. Line plot of disease score (rel FW / symptom category) color coded by symptom category overlaid with scatterplot of CaMV accumulation within the same accession. Identified groups are circled and color coded for their response. Accessions within the circles are written below the graph and named by their accession identifier.

Virus disease in plants is affected by the environment, as well as the genotype of virus and host, the “disease triangle, and manipulation of any edge of this triangle will affect the outcome of virus infection (Hily *et al*., 2016). By controlling for environmental factors in standardized laboratory conditions, as well as for virus genotype by directed infiltration with one strain, we can elucidate the effect of host genotype on CaMV infection in Arabidopsis, through genome-wide association mapping. Previous GWA mappings in Arabidopsis/virus systems have identified resistance loci for the potyvirus Turnip mosaic virus (TuMV), including the well-studied RESTRICTED TEV MOVEMENT 3 (*RTM3*) gene (Rubio *et al*., 2019; Pagny *et al*., 2012; Cosson *et al*., 2010) and novel regulators of RNA silencing during CMV infection (Liu *et al*., 2022). Our GWA mapping identified numerous SNPs associated with differences in CaMV accumulation in agreement with the diverse challenges virus infections impose on host cells. Importantly, no resistance gene is known for Caulimoviruses, except the *CAR1* locus in the Arabidopsis En-2 accession which has not been further mapped (Adhab *et al*., 2018). Nonetheless, several genes involved in various cellular homeostatic processes have been identified through genetic studies that influence CaMV accumulation (Hoffmann *et al*., 2022; Shukla *et al*., 2021; Hafren *et al*., 2017; Schepetilnikov *et al*., 2011; Love *et al*., 2005). Eight out of the 15 T-DNA insertion lines we tested displayed reduced CaMV accumulation in the Col-0 background compared to Col-0 wild type, which appears high as SNPs identified via GWA are frequently effective only in their natural genetic background (Gallois *et al*., 2018; Corwin *et al*., 2016). Notably, none of these eight genes had previously been associated with CaMV disease, underscoring the potential of GWAS to uncover the hidden CaMV disease genes. All of them appear as susceptibility factors for CaMV, as their deletion negatively affects virus accumulation. This could either point to the importance of recessive resistance against CaMV, or more efficient identification of susceptibility factors our GWAS. Even though the identified SNP for *NCED9* has a low allele frequency and was not among the highest scoring ones, the *nced9* mutant had by far the greatest effect and is to our knowledge the most CaMV resistant Arabidopsis T-DNA insertion mutant identified so far. The same T-DNA line has been commonly used and well described for ABA experiments during seed germination, where *NCED9* together with *NCED6* is the main biosynthesis gene (Lefebvre *et al*., 2006; Tan *et al*., 2003b). To our surprise the *nced9* resistance phenotype to CaMV infection and CaMV cannot be rescued by exogenous ABA spray unlike the *aba2-1* mutant. Two other viruses were able to systemically spread through *nced9* and cause wildtype like symptoms, indicating that the resistance is specific for CaMV. While we could not determine yet which function of NCED9 is essential for CaMV infection, the drastic defect in *nced9* mutant warrants more attention.

Interestingly, even though the virus accumulation defect in *nced9* could not be alleviated by exogenous ABA application during CaMV infection, ABA hormone levels had an impact on CaMV accumulation. Plant hormones are an integral part of signaling mechanisms in response to biotic and abiotic environmental stimuli (Verma *et al*., 2016). The level and inducibility of hormonal responses exhibits a large range between Arabidopsis accessions, as identified for the major stress hormones Salicylic acid (Bruessow *et al*., 2021) and ABA (Kalladan *et al*., 2017). Upon pathogen attack, ABA mediates the closure of stomata and deposition of callose at the plasmodesmata to slow the spread of the pathogen (Ton *et al*., 2009). While callose deposition could reduce plasmodesmal trafficking of CaMV as observed for many other viruses (Zavaliev *et al*., 2013; Li *et al*., 2012; Iglesias & Meins, 2000), this is unlikely as spraying with ABA increased systemic CaMV accumulation. Additionally, ABA antagonizes the salicylic acid (SA) mediated systemic required resistance (SAR), which could make it a general target of pathogens to subvert the antimicrobial SAR (Yasuda *et al*., 2008) and could be used by CaMV to escape SAR (Love *et al*., 2005). Yet, for virus infections including CaMV, the role of ABA appears complex. Increased ABA content was measured in *Nicothiana tabacum* after TMV infection (Whenham *et al*., 1986),in rice after *Rice stripe virus* infection (Cui *et al*., 2021), and Cucumber mosaic virus (Alazem *et al*., 2014), but not PPV infection in *Arabidopsis thaliana* (Pasin *et al*., 2020). Further treatment with ABA increased plant resistance to *Tobacco mosaic virus* (Chen *et al*., 2013), *Plum pox virus* (Pasin *et al*., 2020), *Chinese wheat mosaic virus* (He *et al*., 2021) and *Bamboo mosaic virus* (BaMV) in Arabidopsis (Alazem *et al*., 2014) as well as reduced the lesion size in local infections of *Tobacco necrosis virus* (TNV) (Iriti & Faoro, 2008). CaMV on the other hand, accumulated to higher levels upon plant treatment with ABA, in line with a reduction in the *aba2* mutant that was furthermore rescued by ABA application. However, the strong increase in CaMV accumulation upon treatment with the NGDA inhibitor of the ABA biosynthetic NCED family is difficult to understand but notably, ABA and NDGA also acted similarly in that both reduced BaMV accumulation in Arabidopsis (Alazem *et al*., 2014). Thus, our data suggests that CaMV benefits from disruption of ABA homeostasis and indeed, we also found that ABA responsive genes are widely affected by CaMV and highly deregulated when compared to ABA treatment (Hoth *et al*., 2002). This could at least partially be attributable to the CaMV P6 protein interacting with and repressing the function of histone deacetylase H2DC, a regulator of ABA-mediated gene expression (Li *et al*., 2021; Sridha & Wu, 2006).

Taken together, GWA is a powerful tool to identify novel players for DNA virus disease. The large plasticity of Arabidopsis towards CaMV and the independent resistant lines indicate independently evolved resistance mechanism that should be explored further. We found evidence that resistance, as well as tolerance mechanisms play a role during CaMV infection. And lastly, ABA was found as a novel inducer of CaMV accumulation and CaMV infection drastically miss regulates ABA responsive genes.

## Supplementary Data

The following supplementary data are available at JXB online.

Supplementary Dataset S1 contains supplementary Figure S1-S3

Supplementary Figure S1: Additional information in Arabidopsis accessions

Supplementary Figure S2: Correlations between CaMV and CMV data

Supplementary Figure S3: Manhattan plots of GWA-mapping for symptoms and rel. fresh weight

Supplementary Figure S4: Phenotypes of *nced9* and Col-0 plants in different growth conditions

Supplementary Dataset S2 contains supplementary Table S1-S9

Supplementary table S1: Arabidopsis accessions used in this study.

Supplementary table S2: T-DNA lines used in this study with primers used for their confirmation.

Supplementary table S3: Symptom and relative fresh weight data by accessions.

Supplementary table S4: Relative CaMV accumulation in two replicates by accessions.

Supplementary table S5: SNPs that above GWA-score 5 and MAC ≥5.

Supplementary table S6: Genetic elements in 2 kb window of SNPs.

Supplementary table S7: Arabidopsis accessions harboring NCED9 415L.

Supplementary table S8: ABA responsive genes and their expression during CaMV infection.

Supplementary table S9: Primers used in this study.

## Acknowledgements

We’d like to thank Magnus Nordborg for sharing plant material and Thomas Ellis for his insights on GWAS and thoughtful reading of the manuscript. We are thankful to Roger Ling and Andrew Firth for sharing their infectious TRoV clone.

## Author contributions

Conceptualization – GH, AH; Investigation – GH, AS, SL; Formal analysis – GH; AS Visualization – GH; Writing – original draft – GH, AH; Funding acquisition – GH, AH

## Data availability

All data supporting the findings of this study are available within the paper and within its supplementary materials published online.

## References

Adhab, M., Angel, C., Leisner, S. & Schoelz, J.E. (2018). The P1 gene of Cauliflower mosaic virus is responsible for breaking resistance in Arabidopsis thaliana ecotype Enkheim (En-2). Virology, 523, pp. 15–21.

Alazem, M., Lin, K.-Y. & Lin, N.-S. (2014). The abscisic acid pathway has multifaceted effects on the accumulation of Bamboo mosaic virus. Molecular plant-microbe interactions, 27(2), pp. 177–189.

Allen, M., Poggiali, D., Whitaker, K., Marshall, T.R. & Kievit, R.A. (2019). Raincloud plots: a multi-platform tool for robust data visualization. Wellcome open research, 4.

Bartoli, C. & Roux, F. (2017). Genome-Wide Association Studies In Plant Pathosystems: Toward an Ecological Genomics Approach. Frontiers in Plant Science, 8.

Benjamini, Y. & Hochberg, Y. (1995). Controlling the False Discovery Rate: A Practical and Powerful Approach to Multiple Testing. Journal of the Royal Statistical Society: Series B (Methodological), 57(1), pp. 289–300.

Bergès, S.E., Vasseur, F., Bediée, A., Rolland, G., Masclef, D., Dauzat, M., van Munster, M. & Vile, D. (2020). Natural variation of Arabidopsis thaliana responses to Cauliflower mosaic virus infection upon water deficit. PLoS pathogens, 16(5), p. e1008557.

Bergès, S.E., Vile, D., Yvon, M., Masclef, D., Dauzat, M. & van Munster, M. (2021). Water deficit changes the relationships between epidemiological traits of Cauliflower mosaic virus across diverse Arabidopsis thaliana accessions. Scientific Reports, 11(1), p. 24103.

Bruessow, F., Bautor, J., Hoffmann, G., Yildiz, I., Zeier, J. & Parker, J.E. (2021). Natural variation in temperature-modulated immunity uncovers transcription factor bHLH059 as a thermoresponsive regulator in Arabidopsis thaliana. PLOS Genetics, 17(1), p. e1009290.

Butkovic, A., Ellis, T.J., González, R., Jaegle, B., Nordborg, M. & Elena, S.F. (2022). A globally distributed major virus-resistance association in Arabidopsis thaliana. bioRxiv.

Butković, A., González, R., Rivarez, M.P.S. & Elena, S.F. (2021). A genome-wide association study identifies Arabidopsis thaliana genes that contribute to differences in the outcome of infection with two Turnip mosaic potyvirus strains that differ in their evolutionary history and degree of host specialization. Virus Evol, 7(2), p. veab063.

Callaway, A., Liu, W., Andrianov, V., Stenzler, L., Zhao, J., Wettlaufer, S., Jayakumar, P. & Howell, S.H. (1996). Characterization of cauliflower mosaic virus (CaMV) resistance in virus-resistant ecotypes of Arabidopsis. Mol Plant Microbe Interact, 9(9), pp. 810–8.

Cao, J., Schneeberger, K., Ossowski, S., Günther, T., Bender, S., Fitz, J., Koenig, D., Lanz, C., Stegle, O., Lippert, C., Wang, X., Ott, F., Müller, J., Alonso-Blanco, C., Borgwardt, K., Schmid, K.J. & Weigel, D. (2011). Whole-genome sequencing of multiple Arabidopsis thaliana populations. Nat Genet, 43(10), pp. 956–63.

Cecchini, E., Al-Kaff, N.S., Bannister, A., Giannakou, M.E., McCallum, D.G., Maule, A.J., Milner, J.J. & Covey, S.N. (1998). Pathogenic interactions between variants of cauliflower mosaic virus and Arabidopsis thaliana. Journal of Experimental Botany, 49(321), pp. 731–737.

Chen, L., Zhang, L., Li, D., Wang, F. & Yu, D. (2013). WRKY8 transcription factor functions in the TMV-cg defense response by mediating both abscisic acid and ethylene signaling in Arabidopsis. Proceedings of the National Academy of Sciences of the United States of America, 110(21), pp. E1963–E1971.

Chen, P., Lee, B. & Robb, J. (2004). Tolerance to a non-host isolate of Verticillium dahliae in tomato. Physiological and Molecular Plant Pathology, 64(6), pp. 283–291.

Chesnais, Q., Golyaev, V., Velt, A., Rustenholz, C., Brault, V., Pooggin, M.M. & Drucker, M. (2022). Comparative Plant Transcriptome Profiling of Arabidopsis thaliana Col-0 and Camelina sativa var. Celine Infested with Myzus persicae Aphids Acquiring Circulative and Noncirculative Viruses Reveals Virus- and Plant-Specific Alterations Relevant to Aphid Feeding Behavior and Transmission. Microbiol Spectr, 10(4), p. e0013622.

Consortium, G. (2016). 1,135 Genomes Reveal the Global Pattern of Polymorphism in Arabidopsis thaliana. Cell, 166(2), pp. 481–491.

Cooper, J. & Jones, A. (1983). Responses of plants to viruses: proposals for the use of terms. Phytopathology, 73(2), pp. 127–128.

Corwin, J.A., Copeland, D., Feusier, J., Subedy, A., Eshbaugh, R., Palmer, C., Maloof, J. & Kliebenstein, D.J. (2016). The Quantitative Basis of the Arabidopsis Innate Immune System to Endemic Pathogens Depends on Pathogen Genetics. PLoS Genet, 12(2), p. e1005789.

Cosson, P., Sofer, L., Schurdi-Levraud, V. & Revers, F. (2010). A member of a new plant gene family encoding a meprin and TRAF homology (MATH) domain-containing protein is involved in restriction of long distance movement of plant viruses. Plant Signaling & Behavior, 5(10), pp. 1321–1323.

Creelman, R.A., Bell, E. & Mullet, J.E. (1992). Involvement of a Lipoxygenase-Like Enzyme in Abscisic Acid Biosynthesis 1. Plant physiology, 99(3), pp. 1258–1260.

Cui, W., Wang, S., Han, K., Zheng, E., Ji, M., Chen, B., Wang, X., Chen, J. & Yan, F. (2021). Ferredoxin 1 is downregulated by the accumulation of abscisic acid in an ABI5-dependent manner to facilitate rice stripe virus infection in Nicotiana benthamiana and rice. The Plant Journal, 107(4), pp. 1183–1197.

Gallois, J.L., Moury, B. & German-Retana, S. (2018). Role of the Genetic Background in Resistance to Plant Viruses. Int J Mol Sci, 19(10).

Gambetta, G.A., Fei, J., Rost, T.L. & Matthews, M.A. (2007). Leaf scorch symptoms are not correlated with bacterial populations during Pierce’s disease. Journal of Experimental Botany, 58(15-16), pp. 4037–4046.

Gan, X., Stegle, O., Behr, J., Steffen, J.G., Drewe, P., Hildebrand, K.L., Lyngsoe, R., Schultheiss, S.J., Osborne, E.J., Sreedharan, V.T., Kahles, A., Bohnert, R., Jean, G., Derwent, P., Kersey, P., Belfield, E.J., Harberd, N.P., Kemen, E., Toomajian, C., Kover, P.X., Clark, R.M., Rätsch, G. & Mott, R. (2011). Multiple reference genomes and transcriptomes for Arabidopsis thaliana. Nature, 477(7365), pp. 419–423.

Garcia-Ruiz, H., Takeda, A., Chapman, E.J., Sullivan, C.M., Fahlgren, N., Brempelis, K.J. & Carrington, J.C. (2010). Arabidopsis RNA-dependent RNA polymerases and dicer-like proteins in antiviral defense and small interfering RNA biogenesis during Turnip Mosaic Virus infection. Plant Cell, 22(2), pp. 481–96.

González-Guzmán, M., Apostolova, N., Bellés, J.M., Barrero, J.M., Piqueras, P., Ponce, M.a.R., Micol, J.L., Serrano, R.n. & Rodríguez, P.L. (2002). The Short-Chain Alcohol Dehydrogenase ABA2 Catalyzes the Conversion of Xanthoxin to Abscisic Aldehyde[W]. The Plant Cell, 14(8), pp. 1833–1846.

Hafren, A., Macia, J.L., Love, A.J., Milner, J.J., Drucker, M. & Hofius, D. (2017). Selective autophagy limits cauliflower mosaic virus infection by NBR1-mediated targeting of viral capsid protein and particles. Proc Natl Acad Sci U S A, 114(10), pp. E2026–E2035.

Han, S.-Y., Kitahata, N., Sekimata, K., Saito, T., Kobayashi, M., Nakashima, K., Yamaguchi-Shinozaki, K., Shinozaki, K., Yoshida, S. & Asami, T. (2004). A novel inhibitor of 9-cis-epoxycarotenoid dioxygenase in abscisic acid biosynthesis in higher plants. Plant physiology, 135(3), pp. 1574–1582.

Hapiak, M., Li, Y., Agama, K., Swade, S., Okenka, G., Falk, J., Khandekar, S., Raikhy, G., Anderson, A., Pollock, J., Zellner, W., Schoelz, J. & Leisner, S.M. (2008). Cauliflower mosaic virus gene VI product N-terminus contains regions involved in resistance-breakage, self-association and interactions with movement protein. Virus Res, 138(1-2), pp. 119–29.

He, L., Jin, P., Chen, X., Zhang, T.-Y., Zhong, K.-L., Liu, P., Chen, J.-P. & Yang, J. (2021). Comparative proteomic analysis of Nicotiana benthamiana plants under Chinese wheat mosaic virus infection. BMC Plant Biology, 21(1), p. 51.

Hily, J.-M., Poulicard, N., Mora, M.-Á., Pagán, I. & García-Arenal, F. (2016). Environment and host genotype determine the outcome of a plant–virus interaction: from antagonism to mutualism. New Phytologist, 209(2), pp. 812–822.

Hoffmann, G., Mahboubi, A., Bente, H., Garcia, D., Hanson, J. & Hafrén, A. (2022). Arabidopsis RNA processing body components LSM1 and DCP5 aid in the evasion of translational repression during Cauliflower mosaic virus infection. The Plant Cell, 34(8), pp. 3128–3147.

Hoth, S., Morgante, M., Sanchez, J.P., Hanafey, M.K., Tingey, S.V. & Chua, N.H. (2002). Genome-wide gene expression profiling in Arabidopsis thaliana reveals new targets of abscisic acid and largely impaired gene regulation in the abi1-1 mutant. J Cell Sci, 115(Pt 24), pp. 4891–900.

Iglesias, V.A. & Meins, F., Jr. (2000). Movement of plant viruses is delayed in a beta-1,3-glucanase-deficient mutant showing a reduced plasmodesmatal size exclusion limit and enhanced callose deposition. Plant J, 21(2), pp. 157–66.

Iriti, M. & Faoro, F. (2008). Abscisic acid is involved in chitosan-induced resistance to tobacco necrosis virus (TNV). Plant Physiology and Biochemistry, 46(12), pp. 1106–1111.

Kalladan, R., Lasky, J.R., Chang, T.Z., Sharma, S., Juenger, T.E. & Verslues, P.E. (2017). Natural variation identifies genes affecting drought-induced abscisic acid accumulation in <i>Arabidopsis thaliana</i>. Proceedings of the National Academy of Sciences, 114(43), pp. 11536–11541.

Kim, D., Pertea, G., Trapnell, C., Pimentel, H., Kelley, R. & Salzberg, S.L. (2013). TopHat2: accurate alignment of transcriptomes in the presence of insertions, deletions and gene fusions. Genome Biology, 14(4), p. R36.

Kover, P.X. & Schaal, B.A. (2002). Genetic variation for disease resistance and tolerance among Arabidopsis thaliana accessions. Proc Natl Acad Sci U S A, 99(17), pp. 11270–4.

Lefebvre, V., North, H., Frey, A., Sotta, B., Seo, M., Okamoto, M., Nambara, E. & Marion-Poll, A. (2006). Functional analysis of Arabidopsis NCED6 and NCED9 genes indicates that ABA synthesized in the endosperm is involved in the induction of seed dormancy. The Plant Journal, 45(3), pp. 309–319.

Leisner, S.M. & Howell, S.H. (1992). Symptom variation in different Arabidopsis thaliana ecotypes produced by cauliflower mosaic virus. Phytopathology, 82(10), pp. 1042–1046.

Li, H., Handsaker, B., Wysoker, A., Fennell, T., Ruan, J., Homer, N., Marth, G., Abecasis, G., Durbin, R. & Subgroup, G.P.D.P. (2009). The Sequence Alignment/Map format and SAMtools. Bioinformatics, 25(16), pp. 2078–2079.

Li, S., Lyu, S., Liu, Y., Luo, M., Shi, S. & Deng, S. (2021). Cauliflower mosaic virus P6 Dysfunctions Histone Deacetylase HD2C to Promote Virus Infection. Cells, 10(9), p. 2278.

Li, W., Zhao, Y., Liu, C., Yao, G., Wu, S., Hou, C., Zhang, M. & Wang, D. (2012). Callose deposition at plasmodesmata is a critical factor in restricting the cell-to-cell movement of Soybean mosaic virus. Plant Cell Rep, 31(5), pp. 905–16.

Liao, Y., Smyth, G.K. & Shi, W. (2014). featureCounts: an efficient general purpose program for assigning sequence reads to genomic features. Bioinformatics, 30(7), pp. 923–30.

Ling, R., Pate, A.E., Carr, J.P. & Firth, A.E. (2013). An essential fifth coding ORF in the sobemoviruses. Virology, 446(1-2), pp. 397–408.

Liu, S., Chen, M., Li, R., Li, W.-X., Gal-On, A., Jia, Z. & Ding, S.-W. (2022). Identification of positive and negative regulators of antiviral RNA interference in Arabidopsis thaliana. Nature Communications, 13(1), p. 2994.

Long, Q., Rabanal, F.A., Meng, D., Huber, C.D., Farlow, A., Platzer, A., Zhang, Q., Vilhjálmsson, B.J., Korte, A., Nizhynska, V., Voronin, V., Korte, P., Sedman, L., Mandáková, T., Lysak, M.A., Seren, Ü., Hellmann, I. & Nordborg, M. (2013). Massive genomic variation and strong selection in Arabidopsis thaliana lines from Sweden. Nature Genetics, 45(8), pp. 884–890.

Love, A.J., Yun, B.W., Laval, V., Loake, G.J. & Milner, J.J. (2005). Cauliflower mosaic virus, a compatible pathogen of Arabidopsis, engages three distinct defense-signaling pathways and activates rapid systemic generation of reactive oxygen species. Plant Physiol, 139(2), pp. 935–48.

Love, M.I., Huber, W. & Anders, S. (2014). Moderated estimation of fold change and dispersion for RNA-seq data with DESeq2. Genome Biology, 15(12), p. 550.

Martin, M. (2011). Cutadapt removes adapter sequences from high-throughput sequencing reads. 2011, 17(1), p. 3.

Monnot, S., Desaint, H., Mary-Huard, T., Moreau, L., Schurdi-Levraud, V. & Boissot, N. (2021). Deciphering the Genetic Architecture of Plant Virus Resistance by GWAS, State of the Art and Potential Advances. Cells, 10(11), p. 3080.

Montes, N., Alonso-Blanco, C. & Garcia-Arenal, F. (2019). Cucumber mosaic virus infection as a potential selective pressure on Arabidopsis thaliana populations. PLoS pathogens, 15(5), p. e1007810.

Montes, N., Cobos, A., Gil-Valle, M., Caro, E. & Pagán, I. (2021). Arabidopsis thaliana Genes Associated with Cucumber mosaic virus Virulence and Their Link to Virus Seed Transmission. Microorganisms, 9(4), p. 692.

Pagán, I., Alonso-Blanco, C. & García-Arenal, F. (2007). The Relationship of Within-Host Multiplication and Virulence in a Plant-Virus System. PLoS One, 2(8), p. e786.

Pagán, I., Fraile, A., Fernandez-Fueyo, E., Montes, N., Alonso-Blanco, C. & García-Arenal, F. (2010). Arabidopsis thaliana as a model for the study of plant-virus co-evolution. Philosophical transactions of the Royal Society of London. Series B, Biological sciences, 365(1548), pp. 1983–1995.

Pagán, I. & García-Arenal, F. (2020). Tolerance of Plants to Pathogens: A Unifying View. Annu Rev Phytopathol, 58, pp. 77–96.

Pagny, G., Paulstephenraj, P.S., Poque, S., Sicard, O., Cosson, P., Eyquard, J.-P., Caballero, M., Chague, A., Gourdon, G., Negrel, L., Candresse, T., Mariette, S. & Decroocq, V. (2012). Family-based linkage and association mapping reveals novel genes affecting Plum pox virus infection in Arabidopsis thaliana. New Phytologist, 196(3), pp. 873–886.

Pasin, F., Shan, H., García, B., Müller, M., San León, D., Ludman, M., Fresno, D.H., Fátyol, K., Munné-Bosch, S., Rodrigo, G. & García, J.A. (2020). Abscisic Acid Connects Phytohormone Signaling with RNA Metabolic Pathways and Promotes an Antiviral Response that Is Evaded by a Self-Controlled RNA Virus. Plant communications, 1(5), p. 100099.

Paudel, D.B. & Sanfaçon, H. (2018). Exploring the diversity of mechanisms associated with plant tolerance to virus infection. Frontiers in Plant Science, 9, p. 1575.

Prendeville, H.R., Ye, X., Jack Morris, T. & Pilson, D. (2012). Virus infections in wild plant populations are both frequent and often unapparent. American Journal of Botany, 99(6), pp. 1033–1042.

Raybould, A., Maskell, L., Edwards, M.L., Cooper, J. & Gray, A. (1999). The prevalence and spatial distribution of viruses in natural populations of Brassica oleracea. New Phytologist, 141(2), pp. 265–275.

Roossinck, M.J. (2013). Plant virus ecology. PLoS Pathog, 9(5), p. e1003304.

Rubio, B., Cosson, P., Caballero, M., Revers, F., Bergelson, J., Roux, F. & Schurdi-Levraud, V. (2019). Genome-wide association study reveals new loci involved in Arabidopsis thaliana and Turnip mosaic virus (TuMV) interactions in the field. New Phytol, 221(4), pp. 2026–2038.

Schepetilnikov, M., Kobayashi, K., Geldreich, A., Caranta, C., Robaglia, C., Keller, M. & Ryabova, L.A. (2011). Viral factor TAV recruits TOR/S6K1 signalling to activate reinitiation after long ORF translation. The EMBO journal, 30(7), pp. 1343–1356.

Schoelz, J.E. & Leisner, S. (2017). Setting Up Shop: The Formation and Function of the Viral Factories of Cauliflower mosaic virus. Front Plant Sci, 8, p. 1832.

Seren, Ü., Vilhjálmsson, B.J., Horton, M.W., Meng, D., Forai, P., Huang, Y.S., Long, Q., Segura, V. & Nordborg, M. (2012). GWAPP: a web application for genome-wide association mapping in Arabidopsis. The Plant Cell, 24(12), pp. 4793–4805.

Shukla, A., Pagán, I. & García-Arenal, F. (2018). Effective tolerance based on resource reallocation is a virus-specific defence in Arabidopsis thaliana. Mol Plant Pathol, 19(6), pp. 1454–1465.

Shukla, A., Ustun, S. & Hafrén, A. (2021). Proteasome homeostasis is essential for a robust cauliflower mosaic virus infection. bioRxiv.

Soosaar, J.L., Burch-Smith, T.M. & Dinesh-Kumar, S.P. (2005). Mechanisms of plant resistance to viruses. Nature Reviews Microbiology, 3(10), pp. 789–798.

Sridha, S. & Wu, K. (2006). Identification of AtHD2C as a novel regulator of abscisic acid responses in Arabidopsis. Plant J, 46(1), pp. 124–33.

Takahashi, H., Miller, J., Nozaki, Y., Takeda, M., Shah, J., Hase, S., Ikegami, M., Ehara, Y. & Dinesh-Kumar, S.P. (2002). RCY1, an Arabidopsis thaliana RPP8/HRT family resistance gene, conferring resistance to cucumber mosaic virus requires salicylic acid, ethylene and a novel signal transduction mechanism. Plant J, 32(5), pp. 655–67.

Tan, B.-C., Joseph, L.M., Deng, W.-T., Liu, L., Li, Q.-B., Cline, K. & McCarty, D.R. (2003a). Molecular characterization of the Arabidopsis 9-cis epoxycarotenoid dioxygenase gene family. The Plant Journal, 35(1), pp. 44–56.

Tan, B.C., Joseph, L.M., Deng, W.T., Liu, L., Li, Q.B., Cline, K. & McCarty, D.R. (2003b). Molecular characterization of the Arabidopsis 9-cis epoxycarotenoid dioxygenase gene family. Plant J, 35(1), pp. 44–56.

Ton, J., Flors, V. & Mauch-Mani, B. (2009). The multifaceted role of ABA in disease resistance. Trends Plant Sci, 14(6), pp. 310–7.

Torres-Barceló, C., Daròs, J.-A. & Elena, S.F. (2010). HC-Pro hypo-and hypersuppressor mutants: differences in viral siRNA accumulation in vivo and siRNA binding activity in vitro. Archives of virology, 155(2), pp. 251–254.

Verhoeven, A., Kloth, K.J., Kupczok, A., Oymans, G.H., Damen, J., Rijnsburger, K., Jiang, Z., Deelen, C., Sasidharan, R. & van Zanten, M. (2022). Arabidopsis latent virus 1, a comovirus widely spread in Arabidopsis thaliana collections. New Phytologist.

Verma, V., Ravindran, P. & Kumar, P.P. (2016). Plant hormone-mediated regulation of stress responses. BMC Plant Biology, 16(1), p. 86.

Whenham, R., Fraser, R., Brown, L. & Payne, J. (1986). Tobacco-mosaic-virus-induced increase in abscisic-acid concentration in tobacco leaves. Planta, 168(4), pp. 592–598.

Wickham, H. (2016). Package ‘ggplot2’: elegant graphics for data analysis. Springer-Verlag New York. doi, 10, pp. 978–0.

Wickham, H., Averick, M., Bryan, J., Chang, W., McGowan, L., François, R., Grolemund, G., Hayes, A., Henry, L. & Hester, J. Welcome to the tidyverse. J. Open Source Softw. 4 (43), 1686 (2019).

Yasuda, M., Ishikawa, A., Jikumaru, Y., Seki, M., Umezawa, T., Asami, T., Maruyama-Nakashita, A., Kudo, T., Shinozaki, K., Yoshida, S. & Nakashita, H. (2008). Antagonistic interaction between systemic acquired resistance and the abscisic acid-mediated abiotic stress response in Arabidopsis. Plant Cell, 20(6), pp. 1678–92.

Zavaliev, R., Levy, A., Gera, A. & Epel, B.L. (2013). Subcellular dynamics and role of Arabidopsis β-1,3-glucanases in cell-to-cell movement of tobamoviruses. Mol Plant Microbe Interact, 26(9), pp. 1016–30.

